# Direct Measure of DNA Damage in Huntington Disease Reveals Elevated Oxidative Genotoxic Stress and Dysfunctional DNA Repair

**DOI:** 10.1101/2025.11.21.689804

**Authors:** Carlos A. Barba Bazan, Samantha Laville-Dupuy, Maya Klepfish, Jason Cousineau, Tamara Maiuri, Christina Peng, Elizabeth Osterlund, Melanie Alpaugh, Ray Truant

## Abstract

DNA damage repair (DDR) pathway proteins are genetic modifiers of Huntington disease (HD) age of onset and severity. Deficient DDR in HD at the first steps of poly ADP-ribosylation makes DNA damage quantification by traditional downstream DNA damage response markers inaccurate. Repair Assisted Damage Detection (RADD) allows for an accurate direct assessment of oxidative DNA damage, a DDR pathway in which huntingtin protein directly participates. Using RADD, we show that human HD-derived cells have elevated oxidative DNA damage and assess the effect of relevant HD therapeutic targets in rescuing this phenotype. Using huntingtin protein level lowering, we define a dysfunctional role of mutant huntingtin in oxidative DDR. We show that ataxia-telangiectasia mutated (ATM) signaling is deficient and that ATM inhibition rescues elevated oxidative DNA damage in HD cells. In contrast, we show that N6-furfuryladenine (N6FFA) treatment, to increase huntingtin phosphorylation within the amino terminal N17 domain (p-N17), is not effective at restoring HD DDR but reveals dysfunctional N6FFA mediated DDR signaling. We propose a model in which elevated DNA damage arises from both aberrant mutant huntingtin involvement in oxidative DDR and the impairment of oxidative DDR pathways, such as ATM kinase activity, N6FFA processing, and poly ADP-ribose signaling in HD. This highlights the importance of using direct measures of DNA damage such as RADD, rather than measures of a DNA damage response pathway.

## Introduction

Huntington disease (HD) is an autosomal dominant neurodegenerative disorder caused by a CAG trinucleotide repeat expansion in exon 1 of the *HTT* gene translating to a polyglutamine expansion in huntingtin protein (HTT) (1). Although CAG repeat length is inversely correlated with disease onset, considerable variability is present among individuals with the same CAG tract length (2). Genome-wide association studies have identified DNA repair genes as significant modifiers of HD symptom onset, implicating DNA damage repair pathways in disease progression (3).

DNA repair dysfunction is a hallmark of neurodegenerative disorders (4,5), including Alzheimer disease (6), Parkinson disease (7), and amyotrophic lateral sclerosis (8). In HD, nuclear DNA damage has been shown to precede mitochondrial dysfunction in HD peripheral blood mononuclear cells (9), yet the mechanism by which mutant HTT (mHTT) affects DNA repair is unclear. Transcriptional regulation of HTT by p53 suggests that HTT is involved in cellular stress responses (10). Oxidative stress, a common source of DNA damage driven by reactive oxygen species (ROS), has also been heavily implicated in the progression of age-onset neurodegenerative disorders (11–13), likely due to reduced mitochondrial efficiency with ageing. We previously demonstrated that HTT directly participates in DNA repair by scaffolding DNA-repair factors, and phosphorylated N17 HTT (p-N17) accumulates at oxidative DNA damage sites in an ataxia-telangiectasia mutated (ATM)-dependent manner (14). The first 17 amino acids of HTT, termed the N17 domain, have ROS sensing activity (15). The sulfoxidation of a methionine within N17 promotes HTT release from the endoplasmic reticulum enabling subsequent nuclear localization (15–17). N17 phosphorylation at S13 and S16 by casein kinase 2 (CK2) largely regulates this process (18,19). Notably, N17 hypo-phosphorylation has been observed in HD cell models (19–21), and restoring endogenous N17 phosphorylation is protective in HD mouse models (18,21–23).

Considering that DNA repair mechanisms and energy metabolism are dysfunctional in HD, accurately measuring DNA damage is critical (24–27). To directly assess oxidative DNA damage in the context of HD, we employed Repair Assisted Damage Detection (RADD) assays on fibroblasts derived from HD patients with healthy spousal controls, immortalized with human telomerase reverse transcriptase (h-TERT), termed TruHD cells (20). RADD assays use oxidative DNA repair enzymes to label and quantify oxidatively damaged bases (28). This allows for a direct and more sensitive measurement of DNA adducts, due to oxidative stress, rather than γ-H2AX immunofluorescence, a standard DNA damage marker that is increased indirectly as a response to DNA damage. Using RADD, we specifically detected elevated oxidative DNA damage in HD cells, consistent with dysfunctional DNA repair pathways and elevated DNA damage reported in various HD models and patient samples (9,14,29–45).

Poly ADP-ribose (PAR) signaling, one of the earliest steps in DNA damage repair, is deficient in HD, leading to reduced PAR levels in cerebrospinal fluid (CSF) samples from HD patients, even during the prodromal stage of disease (37). Here, we investigate the mechanisms underlying dysfunctional oxidative DNA damage repair in HD, downstream of the PAR response. We provide evidence that ATM signaling is also deficient in HD cells. Upon inhibition of ATM kinase activity, the elevated oxidative DNA damage phenotype in HD cells is rescued, implicating ATM signaling as a potential regulator of HD oxidative DNA repair. Through lowering HTT levels by LMI070 splice modulator, we demonstrate a functional role for HTT in oxidative DNA damage repair that is compromised in HD. Moreover, our attempts to enhance p-N17 levels by N6-furfuryladenine (N6FFA) support that the regulation of p-N17 nuclear localization and signaling is dysfunctional in the oxidative DNA repair response of HD cells. We propose that HTT has an active role in oxidative DNA damage repair that is disrupted by the pathogenic polyglutamine expansion in mHTT. Moreover, we characterize dysfunctional oxidative DNA repair in HD by p-N17 dysregulation and deficient DNA repair pathways, such as N6FFA processing, ATM and PAR signaling.

## Results

### RADD is more sensitive than ɣ-H2AX immunofluorescence in the context of oxidative DNA damage

We have established HTT as a scaffolding protein in the ATM oxidative DNA damage response complex (14). To investigate elevated DNA damage levels driven by deficient DNA repair in HD (9,14,30–35), we used repair assisted DNA damage detection (RADD) assays to directly measure DNA damage (Fig. 1) in hTERT-immortalized HD patient-derived fibroblasts (Table 1), termed TruHD cell lines (20,28). RADD assay relies on enzymatic repair synthesis to label sites of DNA damage. Instead of detecting damage indirectly, RADD exploits the fact that DNA repair enzymes recognize and process damaged bases or strand breaks, creating a site where labeled nucleotides can be incorporated then quantified by immunofluorescence.

**Figure 1:**
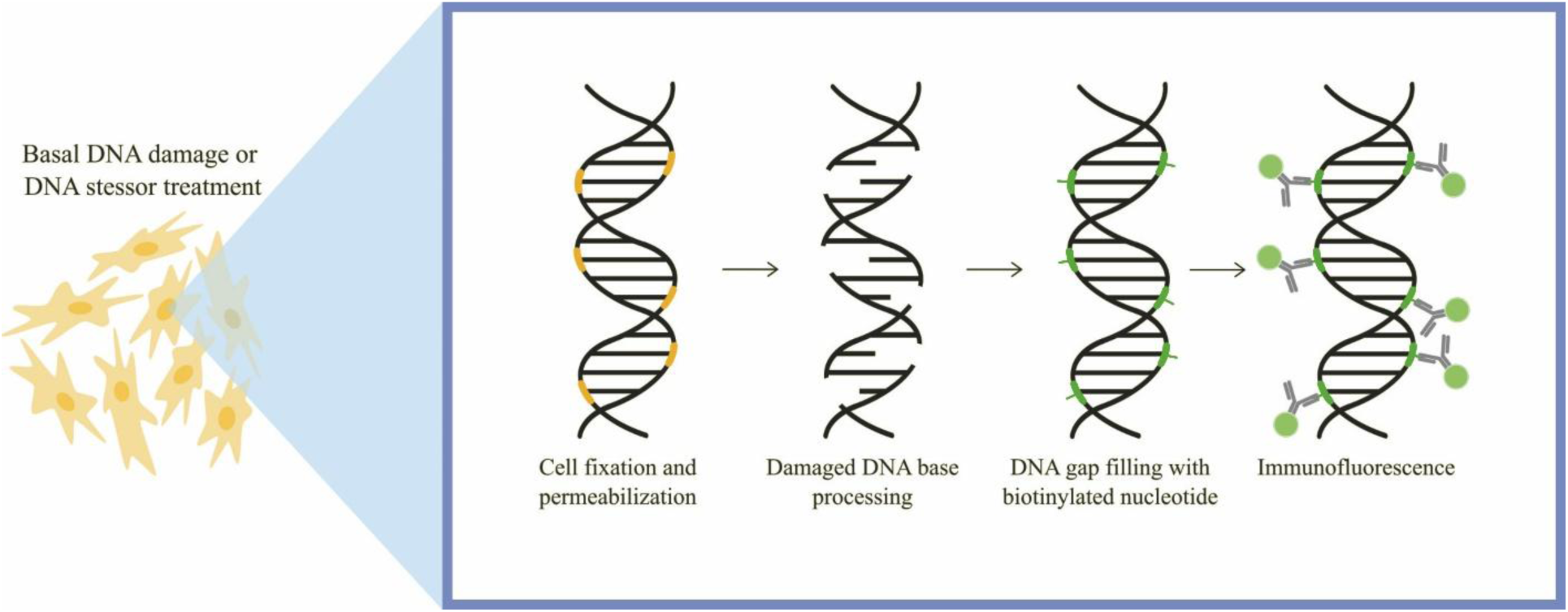
Graphical Abstract of RADD protocol. The RADD protocol starts with the induction of a DNA damage stressor or basal levels of DNA damage. This is followed by fixation and permeabilization of cells. The damaged DNA bases are then modified by an oxidative DNA damage processing enzyme mix followed by the incorporation of biotinylated dUTPs by a DNA gap filling enzyme mix. Cells are then incubated by a primary conjugate antibody against biotin and visualized by widefield fluorescence microscopy.

**Table 1:** Summary of HD cell lines. This table displays CAP (CAG-age product[CAG count – centering constant (30))/scaling constant (6.49)]) score of cells at age of sampling in which scores above 100 indicate expected age of onset (81).

| Primary Cell Name | Cell Line Name | Allele 1 | Allele 2 | Sex | Age at sampling | Cell Type | HD genotype | CAP* Score |
| --- | --- | --- | --- | --- | --- | --- | --- | --- |
| ND30014 | TruHD - Q21Q18F | 21 CAG | 18 CAG | Female | 52 | hTERT-immortalized fibroblasts | Wildtype | N/A |
| ND30013 | TruHD-Q43Q17M | 43 CAG | 17 CAG | Male | 54 | hTERT-immortalized fibroblasts | Heterozygous HD | 108 |
| GM04857 | TruHD-Q50Q40F | 50 CAG | 40 CAG | Female | 23 | hTERT-immortalized fibroblasts | Homozygous HD | 71 |
| GM02185 | TruHD- M36 | Unknown | Unknown | Male | 36 | hTERT-immortalized fibroblasts | Wildtype | N/A |
| GM04767 | TruHD-Q46Q18F | 46 CAG | 18 CAG | Female | 43 | hTERT-immortalized fibroblasts | Heterozygous HD | 106 |
| GM04849 | TruHD-Q52Q45F | 52 CAG | 45 CAG | Female | 28 | hTERT-immortalized fibroblasts | Homozygous HD | 95 |
| NDS00199 | ND38554 | 17 CAG | Unknown | Female | 39 | iPSC-derived neurons | Wildtype | N/A |
| NDS00192 | ND38548 | 42 CAG | Unknown | Female | Unknown | iPSC-derived neurons | Heterozygous HD | N/A |

First, we established RADD as a viable method for detecting oxidative DNA damage in TruHD cells. A subset of RADD assays was performed to directly detect oxidative DNA lesions using bacterial DNA repair enzymes, Fapy-DNA glycosylase, Endonuclease IV and Endonuclease VIII. RADD recognized elevated oxidative DNA lesion levels following treatments with KBrO_3_ and H_2_O_2_, yet not the production of non-oxidative DNA adducts by furfuryl alcohol (Fig. 2A-B). Phosphorylation of H2AX (ɣ-H2AX), which is observed in almost all types of DNA damage (46), is often used as a downstream readout of DNA damage. We compared RADD to ɣ-H2AX immunofluorescence and an internal ROS sensor, 2’,7’-dichlorodihydrofluorescein (H_2_DCFDA), under the same conditions. All three measurements showed increases in respective measurements (Fig. 2A-D). This establishes RADD assays as a viable method to directly investigate oxidative stress in HD.

**Figure 2:**
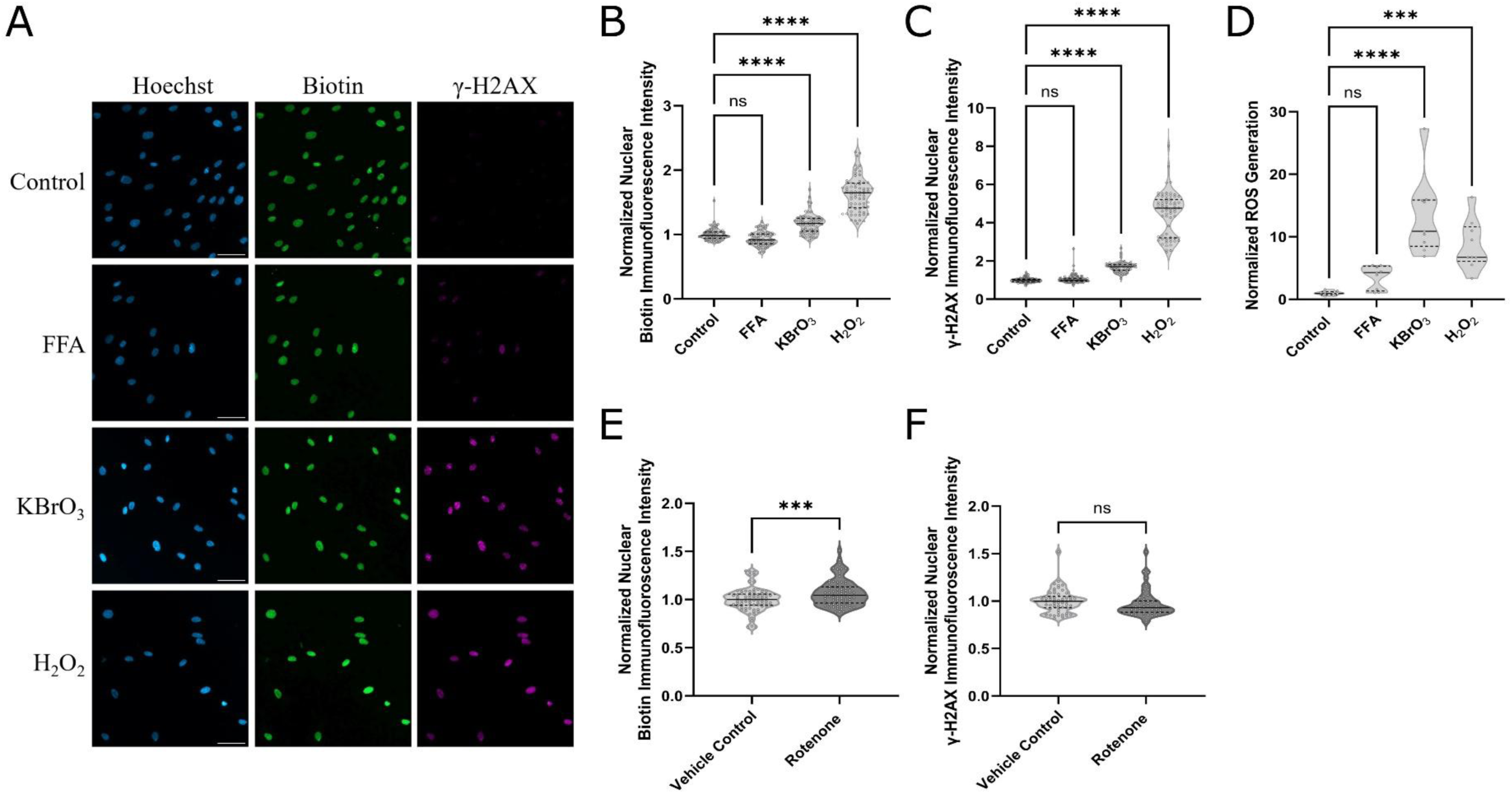
RADD is more sensitive than other measurements of oxidative stress in the context of DNA damage. (A) Representative images of TruHD-Q21Q18 cells showing biotin (RADD) and γ-H2AX nuclear fluorescence intensity. (Scale bars: 50 μm). (B and C) hTERT-immortalized fibroblasts derived from a healthy control (TruHD-Q21Q18) were treated with either 100 mM KBrO_3_, 10 μM H_2_O_2_, or 1 μM FFA for 30 min followed by Repair Assisted Damage Detection (RADD) to detect oxidative DNA lesions (B) and co-stained with an γ-H2AX antibody (C). (D) Comparison of ROS generation by dichlorodihydrofluorescein diacetate (H_2_DCFDA) in TruHD-Q21Q18 cells under indicated oxidative stress treatments as in B and C. (E and F) TruHD-Q21Q18 cells were treated with 1 nM rotenone or vehicle control (DMSO) for 24 hr followed by RADD (E) and co-stained with an γ-H2AX antibody (F). Nuclear RADD and γ-H2AX intensity were measured using CellProfiler, mean nuclear intensity was recorded for each image (18 images per condition; approx. 150 cells per image), and values were normalized to the control conditions. All data is representative of three independent experiments. Statistical analysis was performed using one-way ANOVA with Dunn’s multiple comparisons test (B-D) and unpaired t-test (E-F). Dashed black horizontal lines indicate the interquartile range and medians are indicated by a black horizontal line. (***P < 0.001, ****P < 0.0001).

We next investigated the sensitivity of RADD assays using a mitochondrial complex 1 inhibitor, rotenone, known to increase levels of endogenous ROS (47). At a 1 nM concentration of rotenone for 24hr, RADD assays showed an increase in oxidative DNA lesions while ɣ-H2AX immunofluorescence did not significantly increase (Fig. 2E-F). Thus, RADD assays can more sensitively detect increases in oxidative lesions from genotoxic stressors than markers of strand breaks, such as ɣ-H2AX.

### HD cells have elevated oxidative DNA damage

After validating RADD assays in our cell models, we moved to investigate whether oxidative DNA damage repair is deficient in HD. We have previously shown that HTT interacts with DNA repair proteins in a ROS-dependent manner (14). Since the presence of ROS promotes HTT nuclear localization and recruitment to DNA damage (15,16), we used RADD assays to compare oxidative DNA lesions in TruHD cell lines under steady state conditions and under oxidative stress induced by KBrO_3_. RADD revealed that compared to a healthy control (TruHD-Q21Q18), heterozygous cells carrying one copy of mutant HTT (TruHD-Q43Q17) displayed elevated oxidative DNA damage basally and under conditions of oxidative genotoxic stress (Fig. 3A). Under the same conditions, we assessed ɣ-H2AX immunofluorescence, which detected the increases in oxidative DNA damage with KBrO_3_, but revealed no significant differences between cell lines (Fig. 3B). This further reflects the lower sensitivity of ɣ-H2AX staining (Fig. 2B) and suggests a deficient DNA repair response in HD cells. To further investigate this HD phenotype, we performed RADD on heterozygous and homozygous lines compared to wildtype lines to detect basal oxidative stress. A total of six hTERT-immortalized patient-derived cell lines were assessed and pooled by genotype (Table 1). Heterozygous and homozygous cell lines for the HD mutation displayed elevated oxidative DNA damage compared to wildtype cell lines (Fig. 3C).

**Figure 3:**
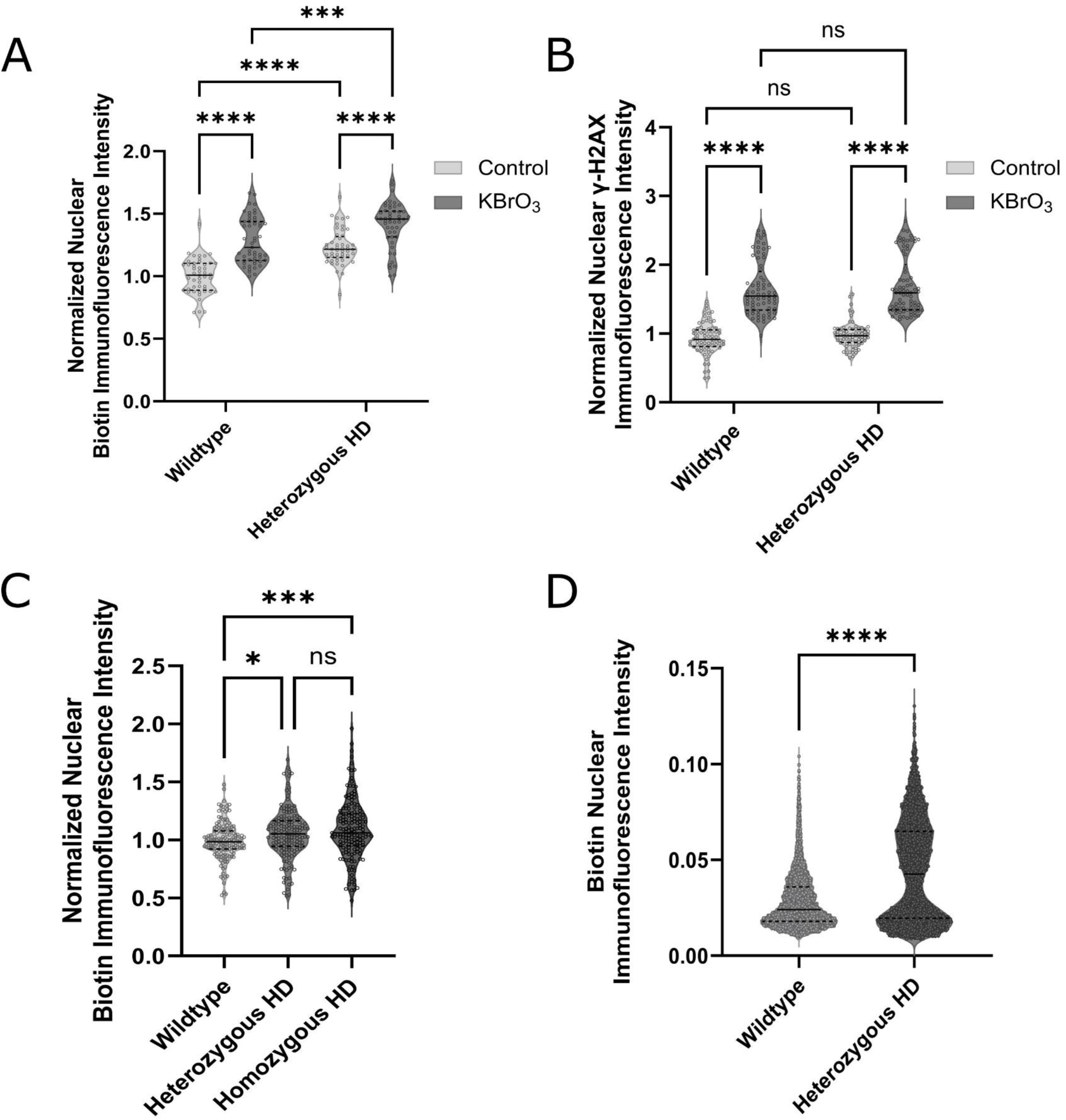
HD cells have elevated oxidative DNA damage levels. (A) hTERT-immortalized fibroblasts derived from a wildtype control (TruHD-Q21Q18) and a heterozygous HD patient (TruHD-Q43Q17) were treated with 100 mM KBrO_3_ for 30 min followed by RADD protocol. (B) TruHD-Q21Q18 and TruHD-Q43Q17 cells were treated with 100 mM KBrO_3_ for 30 min followed by chromatin retention protocol and subsequent staining with a γ-H2AX antibody. (C) hTERT-immortalized fibroblasts were left untreated followed by RADD protocol to detect basal oxidative lesions. Data, separated by genotype, from two wildtype (TruHD-Q21Q18 and TruHD-M36), two heterozygous HD (TruHD-Q43Q17 and TruHD-Q), and two homozygous HD cell lines (TruHD-Q50Q40 and TruHD-Q52Q52) were pooled. Nuclear RADD and γ-H2AX intensity were measured using CellProfiler, mean nuclear intensity was recorded for each image (27 images per condition; approx. 150 cells per image), and values were normalized to the wildtype control conditions. Data is representative of three independent experiments. Statistical analysis was performed using two-way ANOVA with Tukey’s multiple comparisons test (A-B) and one-way ANOVA with Dunn’s multiple comparisons test (C). (D) iPSC-derived neurons were left untreated followed by RADD protocol. Data from two independent differentiations and four technical replicates are shown. Nuclear RADD intensity was measured using CellProfiler, individual cellular nuclear intensity was recorded for each image (n = 400 nuclei were sampled at approx. 10% of total nuclei counts per cell line). Statistical analysis was performed using t-test. Dashed black horizontal lines indicate the interquartile range and medians are indicated by a black horizontal line. (*P < 0.05, ***P < 0.001, ****P < 0.0001).

We then performed RADD on iPSC-derived cortical neurons (Table 1), which also revealed elevated basal oxidative DNA damage levels in HD cells (Fig. 3D). We have previously shown that TruHD-Q43Q17 cells exhibit higher levels of DNA strand breaks than TruHD-Q21Q18 using alkaline comet assays (14). Since the RADD protocol uses enzymes that precisely excise oxidative lesions, these results show that oxidative DNA repair is dysfunctional in HD cells, a human cell model with mutant CAG tracts which does not undergo somatic expansion (20). Thus, dysfunctional oxidative DNA repair is a systemic HD phenotype that does not only affect neurons.

### Huntingtin lowering induces basal oxidative DNA damage in wildtype cells

Given the dysfunctional oxidative DNA repair in HD cells, we next investigated whether reducing HTT expression affects oxidative DNA lesion levels. We used a splice modulator, LMI070, to lower HTT mRNA and protein levels. Due to off-target effects in response to high doses of LMI070 (48,49), we used a low concentration of 1 nM LMI070 for 72 hours. Oxidative DNA lesions were then assessed using RADD under basal and induced oxidative stress conditions. Heterozygous HD cells displayed no significant differences in oxidative damage levels in response to, while wildtype cells showed elevated oxidative DNA lesions with LMI070 treatment in the absence of oxidative stress (Fig 4. A-B). This indicates that a reduction in HTT by LMI070 treatment has no effect on HD cells but leads to dysfunctional basal oxidative DNA repair in wildtype cells. In contrast, RADD assays show a decrease in oxidative DNA lesions in homozygous HD cells with LMI070 treatment and induced oxidative stress (Fig 4. C) This points towards a benefit of decreased mHTT in elevated oxidative stress conditions. This implies that disrupting mHTT involvement in DNA damage repair may be beneficial.

**Figure 4:**
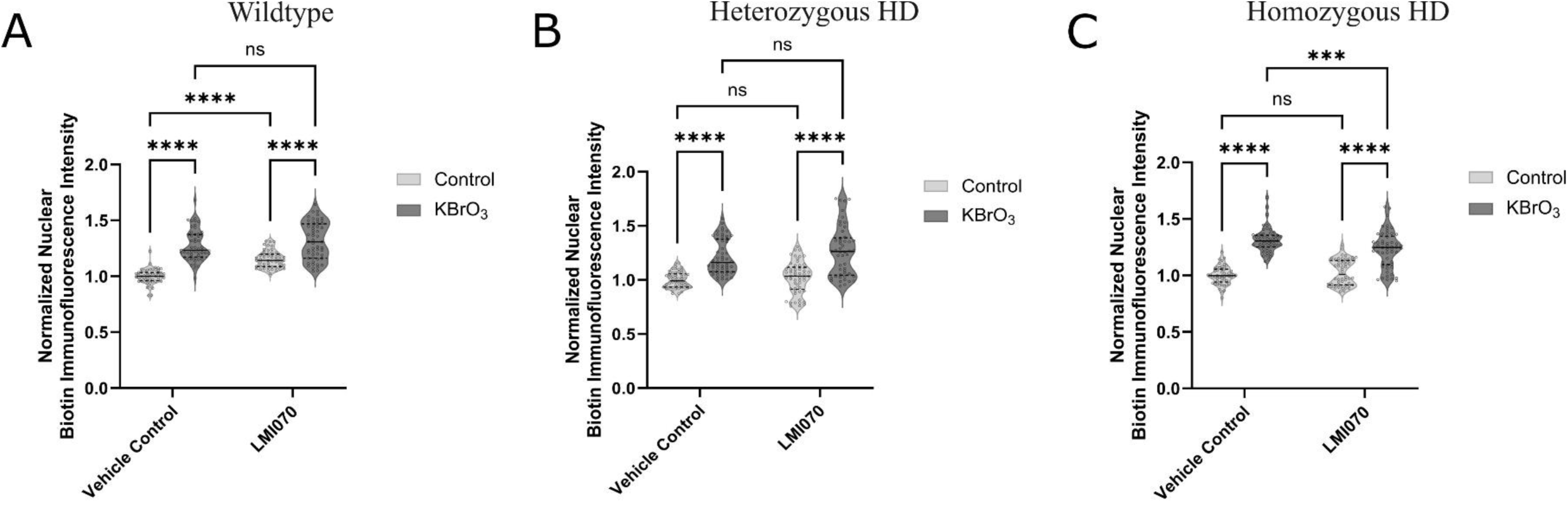
Huntingtin lowering induces basal oxidative DNA damage in wildtype cells. (A-C) hTERT-immortalized fibroblasts derived from a wildtype control (TruHD-Q21Q18) (A), heterozygous HD patient (TruHD-Q43Q17) (B), and homozygous HD patient (TruHD-Q50Q40) (C) were pretreated with 1 nM LMI070 or vehicle control (DMSO) every 24 hr for 72 hr prior to 100 mM KBrO_3_ treatment for 30 min followed by RADD protocol. Nuclear RADD intensity was measured using CellProfiler, mean nuclear intensity was recorded for each image (18 images per condition; approx. 150 cells per image), and values were normalized to the control untreated conditions. All data is representative of three independent experiments. Statistical analysis was performed using two-way ANOVA with Tukey’s multiple comparisons test. Dashed black horizontal lines indicate the interquartile range and medians are indicated by a black horizontal line. (***P < 0.001, ****P < 0.0001).

### ATM inhibition rescues the elevated oxidative DNA damage in HD cells

We next compared how ATM inhibition affects the oxidative stress phenotype in HD cells. We have previously shown that localization of HTT phosphorylated at serines 13 and 16, within N17 to DNA damage happens in an ATM dependent manner (14). ATM is a master regulator in DNA damage repair, which is well characterized in response to double stranded breaks (50). However, ATM is also directly activated in response to ROS (51). Based on ATM inhibition being shown to be neuroprotective in rat striatal neurons (38), we asked whether disrupting HTT localization to damaged DNA affects the elevated DNA damage phenotype in our HD cell models. In the presence of an ATM kinase inhibitor (KU55933), TruHD-Q21Q18 cells displayed no effect while TruHD-Q43Q17 cells showed a significant reduction in elevated RADD phenotype in response to KBrO_3_ (Fig 5A). This suggests that diminishing HTT recruitment to oxidative DNA damage sites by ATM inhibition may be beneficial in HD cells.

**Figure 5:**
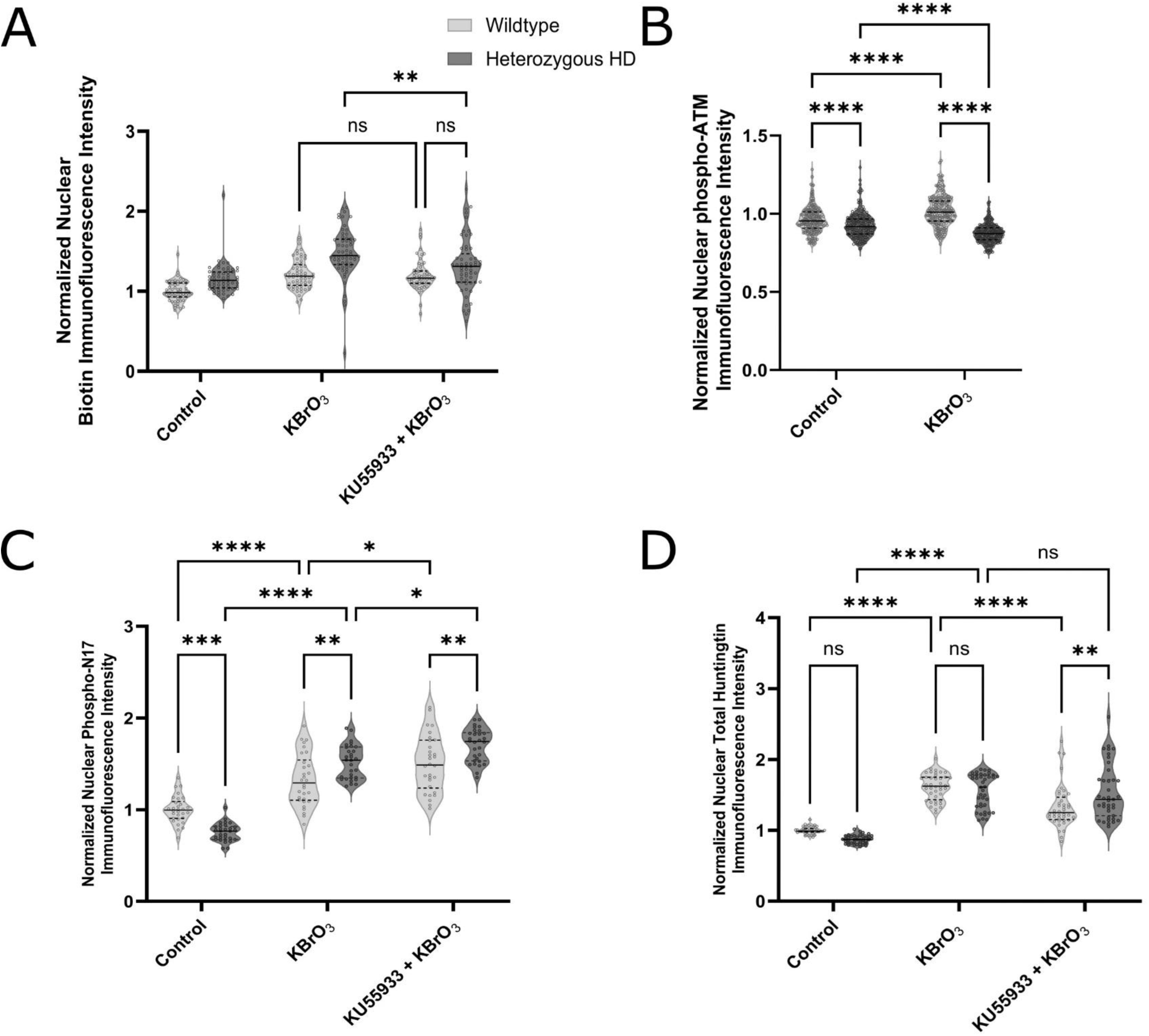
ATM inhibition rescues the elevated oxidative DNA damage in HD cells. (A) hTERT-immortalized fibroblasts derived from a wildtype control (TruHD-Q21Q18) and heterozygous HD patient (TruHD-Q43Q17) were pretreated with 10 μM KU55933 (ATM inhibitor) for 30 min prior to 100 mM KBrO_3_ treatment for 30 min in the presence of KU55933 followed by RADD protocol. Nuclear RADD intensity was measured using CellProfiler, mean nuclear intensity was recorded for each image (18 images per condition; approx. 150 cells per image), and values were normalized to the control untreated conditions. (B) TruHD-Q21Q18 and TruHD-Q43Q17 were treated for 100 mM KBrO_3_ treatment for 30 min followed by chromatin retention protocol and subsequent staining with an ATM(S1981P) antibody. Nuclear ATM intensity was measured using CellProfiler, individual nuclear intensity was recorded, and values were normalized to the control condition (n =400-500 nuclei were sampled at approx. 30% of total nuclei counts per cell line). (C-D) TruHD-Q21Q18 and TruHD-Q43Q17 were pretreated with 10 μM KU5933 (ATM inhibitor) for 30 min prior to treated for 100 mM KBrO_3_ treatment for 30 min followed by chromatin retention protocol and subsequent staining with an antibody against HTT phosphorylated at the N17 domain (phospho-N17) (C) or total HTT (D). Nuclear phospho-N17 and total HTT intensity were measured using CellProfiler, mean nuclear intensity was recorded for each image (10 images per condition; approx. 100 cells per image), and values were normalized to the control untreated conditions. All data is representative of three independent experiments. Statistical analysis was performed using two-way ANOVA with Tukey’s multiple comparisons test. Dashed black horizontal lines indicate the interquartile range and medians are indicated by a black horizontal line. (*P <0.05, **P < 0.01, ***P < 0.001, ****P < 0.0001).

ATM nuclear-cytoplasmic shuttling in response to radiation has been shown to be impaired in HD cell models (34). To examine whether there is dysfunctional ATM signaling in response to oxidative stress in HD cells, we used an antibody against ATM phosphorylated at serine 1981, p-ATM, to measure its nuclear retention. Basal nuclear p-ATM levels were lower in TruHD-Q43Q17 cells compared to TruHD-Q21Q18 cells (Fig 5B). Furthermore, TruHD-Q43Q17 cells do not mount a nuclear p-ATM response to oxidative stress (Fig 5B). Altogether, these data establish a hypo-phosphorylated ATM nuclear response to oxidative stress in HD cells and paradoxically suggests that further impairment of ATM kinase activity is beneficial.

We next compared the localization of p-N17 to the nucleus in response to oxidative stress in TruHD-Q21Q18 and TruHD-Q43Q17 cells. We have previously shown that TruHD-Q43Q17 cells have lower basal whole cell p-N17 levels (20). Immunofluorescence with a phospho-specific antibody against p-N17 revealed that consistent with whole cell levels, basal nuclear p-N17 levels are reduced in HD cells. In contrast, TruHD-Q43Q17 cells show elevated nuclear p-N17 in response to KBrO_3_ compared to TruHD-Q21Q18 cells (Fig. 5C). This indicates that HD cells can mount a p-N17 response to oxidative DNA damage that results in N17 phosphorylation. In addition, inhibition of ATM kinase activity significantly increases p-N17 nuclear staining in both cell lines in response to KBrO_3_ (Fig. 5C).

We then assessed the effect of ATM inhibition on total HTT nuclear retention in response to oxidative DNA damage. In response to KBrO_3_ and ATM kinase inhibition, TruHD-Q21Q18 cells showed a reduction in total nuclear HTT in comparison to treatment with KBrO_3_ alone. Conversely, TruHD-Q43Q17 cells showed no significant differences with ATM kinase inhibition (Fig. 5D). Our findings indicate that there is a disconnect between ATM and HTT in HD oxidative DNA repair. Given that ATM inhibition results in elevated nuclear p-N17, this suggested that increasing p-N17 levels may be beneficial to oxidative DNA repair in HD cells.

### N6FFA regulation of oxidative stress-dependent N17 phosphorylation does not affect HD DNA damage levels

To elucidate how p-N17 levels correlate with oxidative DNA damage repair, we investigated whether modifying N17 phosphorylation affects oxidative DNA damage levels in TruHD cells. Deficiencies in energy metabolism are present in aging brains (52) as well as various neurodegenerative disorders (53). Energy deficits have been well-documented in HD (25–27).

We have previously established that TruHD-Q43Q17 cells have an elevated ADP/ATP ratio (20). CK2, which can utilize kinetin triphosphate, KTP, as a phosphate donor neo substrate, can phosphorylate N17 promoting HTT localization to the nucleus (17–19). We have previously shown that N6-furfuryladenine, or N6FFA, a precursor to KTP, modulates CK2 activity to increase phosphorylation of N17 (18). N6FFA is a natural product of oxidative DNA damage in which adenosine bases undergo the addition of a furfuryl group to generate N6FFA riboside (54). Adenine phosphoribosyltransferase (APRT) salvages N6FFA, excised from the DNA backbone, to eventually generate KTP (55,56).

To assess whether N6FFA affects HTT in the oxidative DNA repair pathway, we treated TruHD cells with N6FFA and measured nuclear retention as measured by p-N17 epitope under basal and oxidative stress conditions. As expected, p-N17 HTT nuclear retention increases with oxidative stress and N6FFA treatment in TruHD-Q21Q18 cells (Fig. 6A). However, p-N17 HTT nuclear retention decreases in TruHD-Q43Q17 cells under the same conditions (Fig. 6B). This points to a p-N17 regulatory function of N6FFA that is dysregulated in HD cells which is further supported by N17 hyperphosphorylation shown in figure 5C. Additionally, basal p-N17 nuclear retention decreases with N6FFA treatment across both cell lines (Fig. 6A-B).

**Figure 6:**
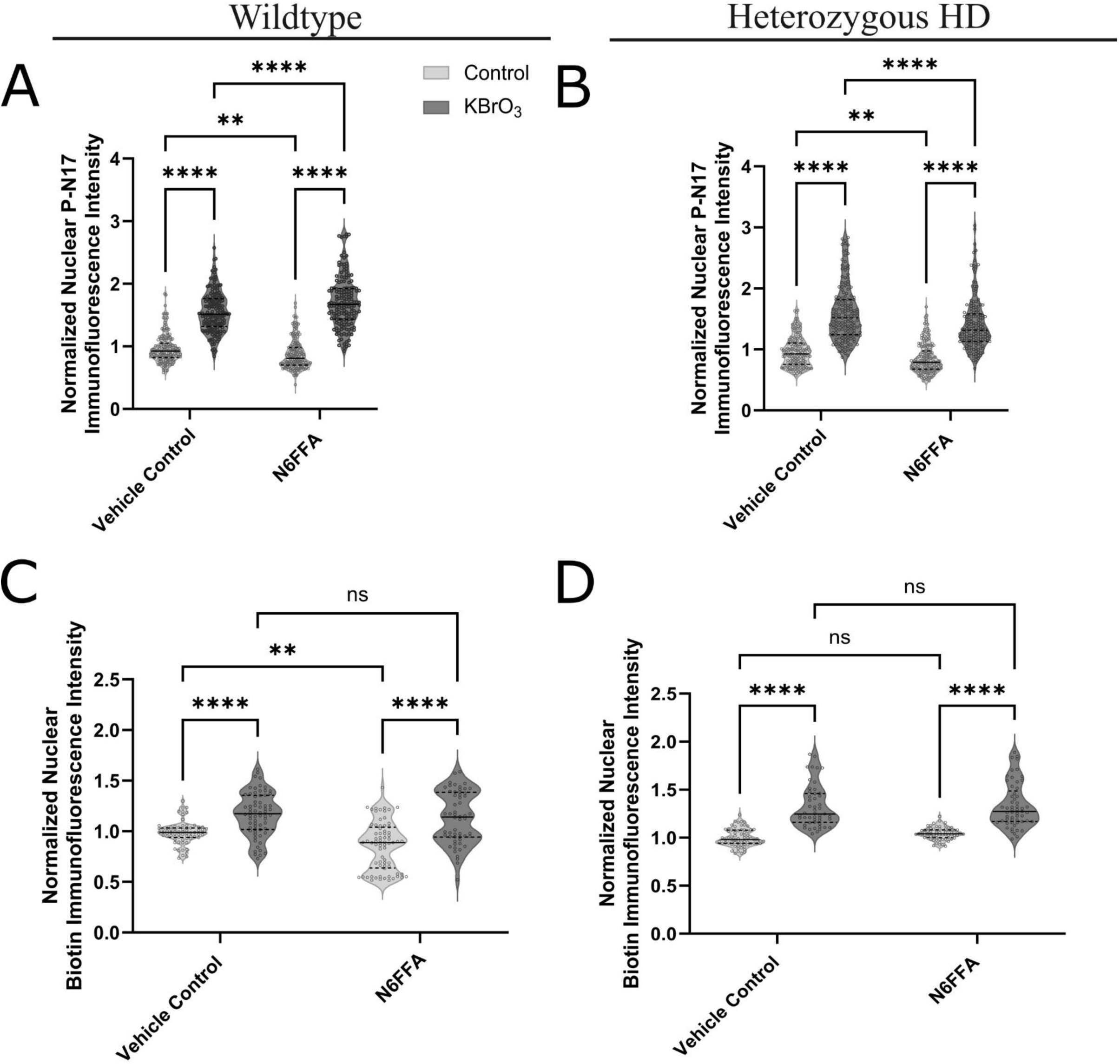
Regulation of oxidative stress-induced N17 phosphorylation by N6FFA does not affect DNA damage levels in HD cells. (A-B) hTERT-immortalized fibroblasts derived from a wildtype control (TruHD-Q21Q18) (A) and heterozygous HD patient (TruHD-Q43Q17) (B) were pretreated with 1 μM N6FFA or vehicle control (NaOH) for 24 hr prior to 100 mM KBrO_3_ treatment for 30 min followed by chromatin retention protocol and subsequent staining with an antibody against HTT phosphorylated at the N17 domain (phospho-N17). Nuclear phospho-N17 intensity was measured using CellProfiler, individual nuclear intensity was recorded, and values were normalized to the untreated control conditions (n=400-500 nuclei were sampled at approx. 30% of total nuclei counts per condition). Data is representative of three independent experiments. (C-E) TruHD-Q21Q18(C), TruHD-Q43Q17 (D) and TruHD-Q50Q40 (E) fibroblasts were pretreated with 1 μM N6FFA or vehicle control (1 μM NaOH) for 24 hr prior to 100 mM KBrO_3_ treatment for 30 min followed by RADD protocol. Nuclear RADD intensity was measured using CellProfiler, mean nuclear intensity was recorded for each image (18 images per condition; approx. 150 cells per image), and values were normalized to the control untreated conditions. Data is representative of four independent experiments. Statistical analysis was performed using two-way ANOVA with Tukey’s multiple comparisons test. Dashed black horizontal lines indicate the interquartile range and medians are indicated by a black horizontal line. (**P < 0.01, ****P < 0.0001).

N6FFA is protective in various HD cell and animal models (18). Therefore, we investigated the effect of N6FFA on oxidative DNA lesions by RADD assays. N6FFA treatment only showed a significant reduction in basal oxidative DNA lesions in TruHD-Q21Q18 cells (Fig 6C). This indicates that elevated chromatin retention of p-N17 under oxidative stress does not result in more effective DNA repair. Additionally, N6FFA treatment did not significantly reduce oxidative DNA lesions in TruHD-Q43Q17 cells (Fig 6D). Since N6FFA treatment only significantly lowers oxidative DNA damage levels in wildtype cells, this points to dysfunctional N6FFA effect on DDR signaling in HD cells.

### Purine metabolism and signaling pathways are deficient in HD

We have previously shown that p-N17, CK2, APRT and N6FFA riboside all accumulate at induced DNA damage sites (18). This suggests that all the components of the N6FFA salvaging pathway facilitate N17 phosphorylation. To further investigate N6FFA processing in HD, we compared the naturally occurring N6FFA riboside levels in the chromatin of TruHD cells using immunofluorescence. Basally, TruHD-Q43Q17 cells were found to have elevated N6FFA riboside levels compared to TruHD-Q21Q18 cells (Fig. 7A). This suggests HD cells have dysfunctional N6FFA riboside excision. Further, figure 6B implies that HD cells also have dysfunctional downstream processing of N6FFA in N17 phosphorylation.

**Figure 7:**
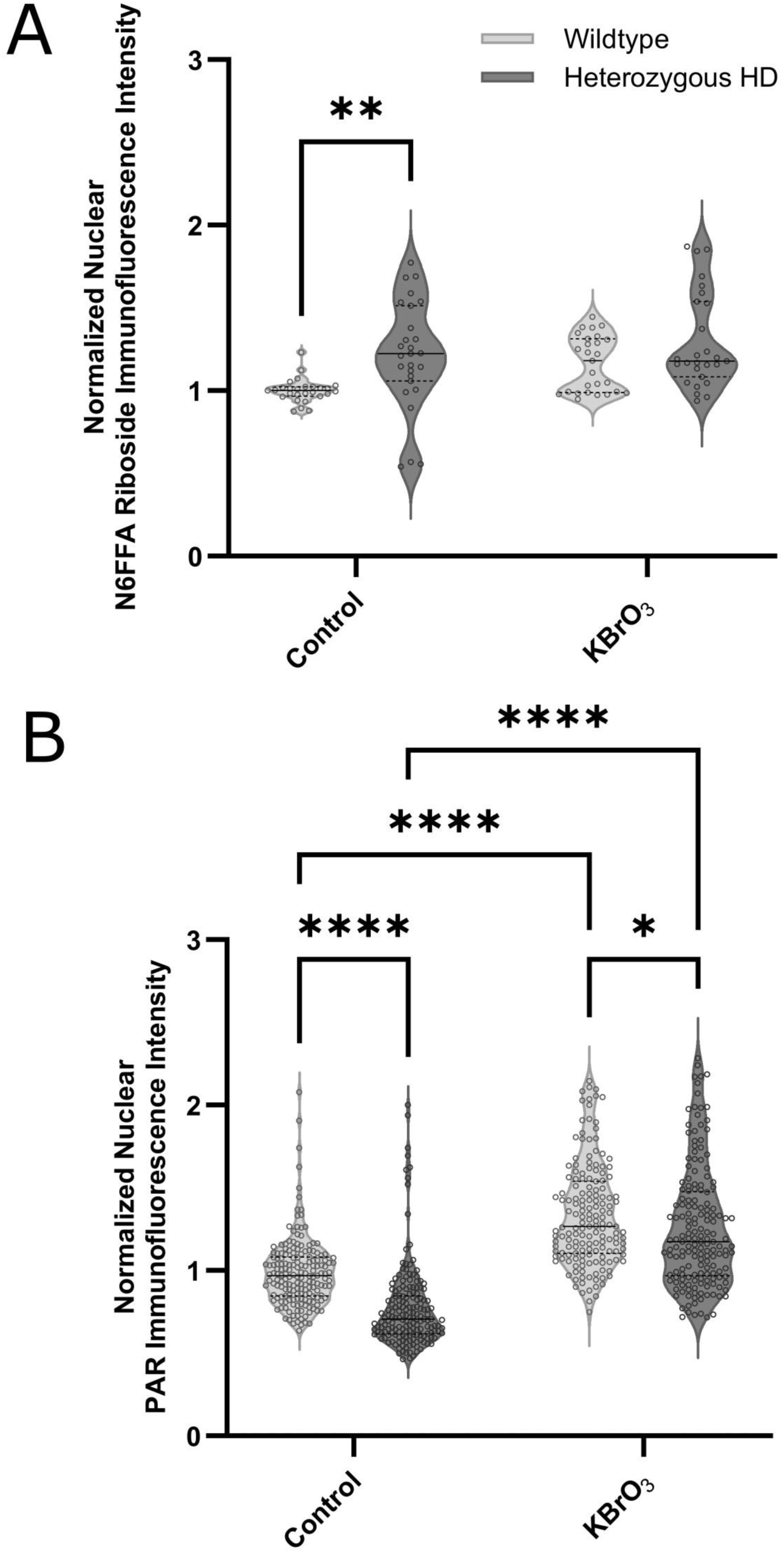
Purine metabolism and signaling pathways are deficient in HD. (A) hTERT-immortalized fibroblasts derived from a wildtype control (TruHD-Q21Q18) and heterozygous HD patient (TruHD-Q43Q17) were treated with 100 mM KBrO_3_ treatment for 30 min followed by antigen presentation protocol and subsequent staining with an antibody against N6FFA riboside. Nuclear N6FFA riboside was measured using CellProfiler, mean nuclear intensity was recorded for each image (9 images per condition; approx. 150 cells per image), and values were normalized to the control untreated conditions. (B) TruHD-Q21Q18 and TruHD-Q43Q17 were treated with 100 mM KBrO_3_ treatment for 30 min followed by methanol fixation and subsequent staining with a poly ADP-ribose (PAR) antibody. Nuclear PAR intensity was measured using CellProfiler, individual nuclear intensity was recorded, and values were normalized to the control condition (n=400-500 nuclei were sampled at approx. 30% of total nuclei counts per cell line). Data is representative of three independent experiments. Statistical analyses were performed using two-way ANOVA with Tukey’s multiple comparisons test. Dashed black horizontal lines indicate the interquartile range and medians are indicated by a black horizontal line. (*P<0.05, **P<0.01, ****P < 0.0001).

Our results suggest that dysfunctional N6FFA signaling may contribute to energy deficits in HD. We previously established PAR signaling, an energy expensive process, is impaired in HD patients and TruHD lines (37). We showed that when co-staining HD fibroblasts for RADD and PAR, DNA damage levels were increased in HD cells while PAR levels were not significantly different than wild type cells, indicating an impaired PAR response. Here, using a staining method without involving chromatin retention, we can better capture the PAR deficiency and observe that TruHD-Q43Q17 cells have deficient PAR signaling basally and in response to oxidative stress compared to TruHD-Q21Q18 cells (Fig. 7B), corroborating our previous cerebral spinal fluid data from HD patients and TruHD cells (37). Deficient PAR signaling and N6FFA processing can be attributed to the dysfunctional purinergic pathways in HD (57,58).

## Discussion

Current DNA adduct quantification methods are limited by the lack of specificity in DNA damage repair pathway analysis, reliance on secondary readouts, or the introduction of further DNA damage. Here, we used RADD assays, to specifically detect oxidative DNA lesions, to investigate the effect of HTT lowering, ATM inhibition and N6FFA treatment in patient-derived hTERT-immortalized fibroblasts. Ɣ-H2AX is often used as a downstream marker of many types of DNA damage (46), however, it relies on an intact pathway response to DNA damage. We showed that RADD assays are more sensitive than ɣ-H2AX immunofluorescence at identifying oxidative DNA repair dysfunction. By RADD, we reveal elevated oxidative DNA damage in multiple HD cell models. Additionally, our findings highlight the importance of considering energy deficits in HD and other neurodegenerative disorders when investigating DNA damage repair. Direct measurement of DNA damage adducts in specific pathways via RADD should be prioritized rather than indirect measures of DNA damage repair signaling processes that require energy, such as the H2AX phosphorylation and PARylation, and hence are inhibited in HD. This is supported by conflicting data on ɣ-H2AX signaling reported in various HD cell models (33,34,38,59,60).

### Mutant HTT has both loss and gain of toxic function in oxidative DNA damage repair

We sought to investigate how mHTT alters the oxidative DNA damage response using RADD assays. Basally, HTT lowering by LMI070 induced oxidative DNA lesions in wildtype cells while HD cells were not affected. This points to the role of HTT in oxidative DNA damage repair while displaying the compromised function of mHTT. Our data suggests that mHTT at DNA damage sites is not the only driver of oxidative DNA repair dysfunction. Only homozygous HD cells displayed a reduction in oxidative DNA lesions with induced oxidative stress under LMI070 treatment. Together, these data suggest that mHTT loses function in basal oxidative DNA damage repair while concurrently being detrimental to induced oxidative DNA damage repair. This is consistent with a toxic gain of function of mHTT in the context of age-onset ROS stress.

### Dysregulated ATM signaling contributes to the elevated oxidative DNA repair phenotype in HD cells

ATM signaling has been implicated in multiple neurodegenerative disorders. An autosomal recessive loss-of-function mutation in the *ATM* gene is the cause of ataxia-telangiectasia (A-T) (61). ATM kinase activity has also been shown to regulate the localization of the ataxin-1 protein to DNA damage, which is affected in spinocerebellar ataxia type 1, another polyglutamine expansion disorder (62). For HD, we have previously shown that ATM kinase activity also regulates nuclear HTT localization to DNA damage sites (14). Furthermore, ATM signaling has been shown to be hyperactive in HD cells (38). However, these findings either rely on measuring ATM phosphorylation targets, such as ɣ-H2AX, or do not directly investigate the nuclear response of ATM to DNA damage (33,38).

We directly analyzed the chromatin retention of ATM and HTT as opposed to their nuclear recruitment to investigate the dysfunctional oxidative DNA damage response in HD. Using this method, we show that HD cells experience deficient ATM signaling in the oxidative DNA repair response. We also show that ATM inhibition under oxidative stress only reduces total nuclear HTT in wildtype cells, further pointing to dysfunctional ATM signaling in HD cells. This is supported by deficient radiation-induced nuclear-cytoplasmic shuttling of ATM in HD cell models (34). Our results could explain the reduced ATM-mediated 53BP1 puncta formation in response to DNA damage and increased topoisomerase I cleavage complexes which are associated with ATM deficiency, previously observed in HD cells (41). Additionally, deficient ATM signaling could contribute to classic HD phenotypes such as elevated ROS levels, impaired mitochondrial function, and Purkinje cell degeneration (63,64), which are also characteristic of ataxia telangiectasia (65–69).

In addition to deficient ATM signaling, we paradoxically observed the rescue of the elevated oxidative DNA damage phenotype in HD cells with ATM inhibition. This infers that elevated DNA damage stems from mHTT involvement in ATM-mediated DNA repair. An explanation comes from our previous findings showing the loss of PARP1-stimulating activity by mHTT and deficient PAR signaling in HD patient CSF and cell samples (37). Deficient PAR signaling elevates ATM signaling and processing of subsequent DNA damage (70–72). Notably, we confirmed that our HD cell model, TruHD-Q43Q17, also has a deficient PAR response which translates to clinical observations in HD patient CSF samples.

We propose that mHTT impairs PAR signaling, ATM activity and their coordinated functions in the DNA damage response in HD. Thus, the rescue of dysfunctional DNA repair by ATM inhibition in HD extends beyond mHTT localization to DNA damage sites but also interferes with ATM-dependent DNA repair pathways. While ATM inhibition has been shown to be protective in HD cell models (38), this may not be a viable therapeutic approach. ATM is critical for genomic stability and is a key regulator of DNA damage repair, thus long-term ATM inhibition could have detrimental effects in HD patients, which already experience deficient PAR signaling (37).

### Dysfunctional N6FFA processing leads to dysregulated N17 phosphorylation in HD

Elevated basal N6FFA riboside in HD cells suggests there is dysfunctional N6FFA excision, although little is known about N6FFA metabolism in DNA repair. However, impaired purinergic pathways have been noted in HD (57,58). Our data on deficient PAR signaling and N6FFA processing expand on dysfunctional purine metabolism and signaling pathways in HD. Treatment with N6FFA, which modulates nuclear N17 phosphorylation by CK2 in an oxidative stress dependent manner (18), reduced basal oxidative DNA damage in wildtype cells, with no effect in HD cells, consistent with lower nuclear p-N17. The protective effect of N6FFA against ROS in human cells has been well established (73–78). The absence of this protective effect in HD cells suggests that N6FFA utilization in signaling is disrupted by mHTT. Therefore, the protective effect of N6FFA in HD models may stem from regulating the ability of CK2 to phosphorylate p-N17 and other substrates.

Our RADD data, based on ATM inhibition and N6FFA treatment, indicate that nuclear N17 phosphorylation does not directly correlate with oxidative DNA damage levels. Furthermore, dysfunctional DNA repair in HD is driven by mHTT interference in oxidative DNA damage repair pathways, rather than solely from its localization at DNA damage sites. Overall, our observations reveal a consistent disconnect between mHTT and oxidative DNA repair signaling pathways. Understanding this relationship is critical for identifying effective therapeutic targets for HD and gaining mechanistic insights into neurodegeneration.

## Materials and Methods

### Reagents

All reagents for RADD were purchased from New England Biolabs unless otherwise stated.

### Antibodies

Rabbit polyclonal antibodies against phosphorylated serines 13 and 16 in the N17 domain of HTT (1:500, anti-p-N17, New England Peptides) were previously developed and validated (19,20). Additional antibodies used in this study were: mouse monoclonal anti-p-ATM (1:200, 10H11.E12, Santa Cruz Biotechnology), rabbit PAR (1:500, MABE1031, MilliporeSigma) detection reagent, rabbit monoclonal anti-HTT (1:500, EPR5526, Abcam), rabbit monoclonal anti-ɣH2AX (1:250, ab81299, Abcam), rabbit polyclonal anti-N6FFA riboside (1:100, AS09 443, Agrisera), mouse monoclonal Alexa Fluor 488-conjugated anti-biotin (1:500, BK-1/39, ThermoFisher), secondary antibodies against rabbit and mouse IgG (1:500, Alexa Fluor 488/Alexa Fluor 594, Abcam).

### Cell Culture

#### TruHD cells

Patient fibroblasts (ND30014, ND30013, GM04857, GM02185, GM04767, GM04849) were purchased from Coriell Institute. hTERT-immortalization of these fibroblasts, TruHD cells, was previously described (20). Cells were cultured in MEM (Life Technologies #10370) with 15% fetal bovine serum (FBS, Life Technologies) and 1X GlutaMAX (Life Technologies #35050) and grown at 37°C with 5% CO_2_ and 8% O_2_.

#### Neuronal Differentiation of iPSCs

iPSCs (NDS00199, NDS00192) were purchased from NIH and differentiated into neurons as previously described (79). Samples were unblinded after analysis.

### Treatments

#### Induction of DNA damage

100 mM potassium bromate (KBrO_3_), 1 μM furfuryl alcohol (FFA) and 10 μM hydrogen peroxide (H_2_O_2_) were diluted in phosphate-buffered saline (PBS, supplemented with Ca^2+^ and Mg^2+^). Rotenone was diluted in DMSO to 1 nM for inhibition of mitochondrial complex 1.

#### Huntingtin lowering

Cells were pre-treated with 1 nM LMI070 (Selleckchem), diluted in DMSO, in media every 24 hr for 72 hr, prior to KBrO_3_ treatments.

#### ATM inhibition

Cells were pre-treated with 10 μM ATM inhibitor (KU55933, Selleckchem), diluted in DMSO, in PBS (supplemented with Ca^2+^ and Mg^2+^) for 30 min with subsequent KBrO_3_ treatments including 10 μM KU55933.

#### N6FFA treatments

Cells were pre-treated with 1 μM N6FFA, diluted in NaOH, in media for 24hr, prior to KBrO_3_ treatments.

### Immunofluorescence

#### Chromatin Retention

TruHD cells were seeded in glass-bottom 8-well plates (Cellvis) to ∼95% confluence. After treatments, cells were washed with PBS (supplemented with Ca^2+^ and Mg^2+^). Soluble proteins were extracted with 0.2% Triton X-100 in PBS for 2 min on ice. Next, cells were PBS washed and fixed with methanol for 15 min at -20°C. After PBS washing, cells were incubated with blocking buffer (10% FBS in PBS) for 30 min at room temperature. Cells were incubated with primary antibodies, diluted in blocking buffer for 1 hr at room temperature or overnight at 4°C, followed by PBS washes. Then, cells were incubated with secondary antibodies, diluted in blocking buffer, for 30 min at room temperature. After PBS washing, nuclei were stained with Hoechst (0.2 μg/mL in PBS) for 4 min at room temperature, followed by PBS washes and imaging in PBS using a 20× objective on the EVOS FL Auto 2 widefield microscope. Nuclei were identified by Hoechst and nuclear intensity of immunostaining was measured using CellProfiler.

#### Methanol fixation

TruHD cells were seeded in glass-bottom 8-well plates (Cellvis) to ∼95% confluence. After treatments, cells were washed with PBS (supplemented with Ca^2+^ and Mg^2+^), fixed with methanol for 15 min at -20°C. Protocol then follows the subsequent steps in chromatin retention.

#### Antigen presentation

TruHD cells were seeded in glass-bottom 8-well plates (Cellvis) to ∼95% confluence. After treatments, cells were washed with PBS (supplemented with Ca^2+^ and Mg^2+^) and fixed with 1:1 methanol-acetone for 10 min at -20°C. Next, cells are hydrated with PBS for 5 min at room temperature, followed by an incubation with 2N HCl for 45 min at room temperature. After PBS washes, cells are neutralized with 50 mM Tris-HCl pH 8.8 for 5 min at room temperature, followed by PBS washing. Protocol then follows blocking buffer incubation and subsequent steps in chromatin retention.

### RADD assay

Cells were seeded in glass-bottom 8-well, 96-well or 24-well plates (Cellvis) to ∼95% confluence. Cells were washed with PBS and then incubated with either PBS (Control) or treatments (100 mM KBrO_3_, 1 μM FFA, or 10 μM H_2_O_2_), diluted in PBS, for 30 minutes. Alternatively, seeded cells were incubated with 0 nM or 1nM rotenone, diluted in DMSO, in media for 24 hr at 37°C with 5% CO_2_ and 5 to 8% O_2_. After PBS washing, RADD assays were performed as previously described (26). Cells were imaged in PBS using 20× objectives on the EVOS FL Auto 2 widefield microscope or Zeiss Cell Discover 7. Nuclei were identified by Hoechst and nuclear biotin intensity was measured using CellProfiler.

### Reactive oxygen species generation measurement

The generation of ROS in TruHD cells was detected using 2′,7′-dichlorodihydrofluorescein diacetate (H_2_DCFDA). Reactive oxygen species generation assay was performed as previously described (80).

### Statistical Analysis

All Data are represented as a mean with three independent trials unless otherwise specified in figure captions. Statistical analysis was performed using GraphPad Prism V and specific analyses are described in figure captions for each experiment.

## Acknowledgements

We thank Dr. Natalie Gassman for her insight into applying RADD to our Huntington disease cell models.

## Abbreviations

| Abbreviation | Definition |
| --- | --- |
| DNA | Deoxyribonucleic acid |
| DDR | DNA damage repair |
| HD | Huntington disease |
| RADD | Repair assisted damage detection |
| ATM | Ataxia-telangiectasia mutated |
| N6FFA | N6-furfuryladenine |
| p-N17 | N17 domain of huntingtin phosphorylated at serines 13 and 16 |
| HTT | Huntingtin |
| mHTT | Mutant huntingtin |
| ROS | Reactive oxygen species |
| CK2 | Casein kinase 2 |
| h-TERT | Human telomerase reverse transcriptase |
| γ-H2AX | Phosphorylated H2A histone variant |
| PAR | Poly ADP-ribose |
| CSF | Cerebrospinal fluid |
| H<sub>2</sub>DCFDA | 2',7'-dichlorodihydrofluorescein |
| iPSC | Induced pluripotent stem cell |
| ADP | Adenosine diphosphate |
| ATP | Adenosine triphosphate |
| KTP | Kinetin triphosphate |
| APRT | Adenine phosphoribosyltransferase |
| BBC3 | BCL2 binding component 3 |
| A-T | Ataxia-telangiectasia |
| PARP | Poly ADP-ribose polymerase |
| ATXN3 | Ataxin 3 |
| PNKP | Polynucleotide kinase 3'-phosphatase |
| XRCC1 | X-ray repair cross complementing 1 |
| 53BP1 | P53 binding protein 1 |
| A2AR | Adenosine A2A receptor |
| KBrO<sub>3</sub> | Potassium bromate |
| FFA | Furfuryl alcohol |
| H<sub>2</sub>O<sub>2</sub> | Hydrogen peroxide |
| PBS | Phosphate-buffered saline |
| DMSO | Dimethyl sulfoxide |
| FBS | Fetal bovine serum |
| HCl | Hydrochloric acid |
| CAP | CAG-age product |

